# Low-cost microphysiological systems: Feasibility study of a tape-based barrier-on-chip system for small intestine modeling

**DOI:** 10.1101/2020.01.06.894147

**Authors:** Thomas E. Winkler, Michael Feil, Eva F.G.J. Stronkman, Isabelle Matthiesen, Anna Herland

## Abstract

We see affordability as a key challenge in making organs-on-chips accessible to a wider range of users, particularly outside the highest-resource environments. Here, we present an approach to barrier-on-a-chip fabrication based on double-sided pressure-sensitive adhesive tape and off-the-shelf polycarbonate. Besides a low materials cost, common also to PDMS or thermoplastics, it requires minimal (€ 100) investment in laboratory equipment, yet at the same time is suitable for upscaling to industrial roll-to-roll manufacture. We evaluate our microhpysiological system with an epithelial (C2BBe1) barrier model of the small intestine, studying the biological effects of permeable support pore size, as well as stimulation with a common food compound (chili pepper-derived capsaicinoids). The cells form tight and continuous barrier layers inside our systems, with comparable permeability but superior epithelial polarization compared to Transwell culture, in line with other perfused microphysiological models. Permeable support pore size is shown to weakly impact barrier layer integrity as well as the metabolic cell profile. Capsaicinoid response proves distinct between culture systems, but we show that impacted metabolic pathways are partly conserved, and that cytoskeletal changes align with previous studies. Overall, our tape-based microphysiolgical system proves to be a robust and reproducible approach to studying physiological barriers, in spite of its low cost.

## Introduction

Microphysiological systems have the potential to reduce animal testing, to accelerate drug development, and to study cellular processes that are simply not accessible in live humans.^[1]^ To date, however, high costs represent a significant barrier to entry into the field (Table 1). This applies, on the one hand, to commercial solutions. Companies like Emulate or TissUse have surmounted scalability challenges, yet costs remain high. It applies, on the other hand, also to in-house fabrication of organs-on-chips in an academic setting. Set-up even for the predominantly-used poly(dimethylsiloxane) (PDMS)-based processing can be prohibitive for researchers in low-resource environment. Indeed, research from the top ten countries in terms of per-capita research spending accounts for 78% of the primary literature on organs-on-chips (compared to only 51% for broadly-defined *in-vitro* research, or 59% for labs-on-chips).^[2,3]^

**Table 1:** Overview of Organ-on-Chip Approaches & Costs. * 40 “organs” for simultaneous use per plate, a format somewhat more suited to high-throughput testing than academic research. It is also worth noting that the Mimetas approach uniquely eliminates the need for investment in an expensive pump or control system, but the lack of true perfusion also carries inherent limitations.

|  | Tape<br>(this work) | PDMS | Thermo-<br>plastics |
| --- | --- | --- | --- |
| Base materials<br>cost / “organ” | € 0.5 | € 0.3 | € 0.2 |
| Tooling cost | € 0 | € 100+ | € 1,000+ |
| Equipment cost | € 100-1,000+ | € 1,000-10,000+ | € 5,000-50,000+ |
| Assembly aides | Pressure only | Plasma, Chemical | Heat+Pressure,<br>Chemical |
| Scalability | Roll-to-roll | Difficult | Inherently |
| Commercial<br>example | None | Emulate Inc (US)<br>>€ 100/“organ” | Mimetas BV (NL)<br>>€ 350/unit* |

We see the current high-resource fabrication requirements for organs-on-chips as the key barrier to “democratization” of the field. Unlike labs-on-chips, microphysiological systems almost exclusively require integration of disparate materials – all capable of supporting cell culture – to create biomimetic compartmentalization. Prime examples are epithelial or endothelial barrier models, which make up the largest fraction of modeled organs.^[4]^ One review paper for blood-brain-barrier devices – exemplary of the larger field – shows that fabrication is dominated by combinations of glass, PDMS, and thermoplastics.^[5]^ Even the simplest bonding scenario of glass/PDMS needs to be facilitated by external means (plasma), requiring additional equipment and making scale-up challenging.

Double-coated pressure-sensitive adhesive tape (hereafter, tape) offers an ideal solution to the challenges described. First, it is a very affordable material that can be patterned with minimal cost (a scalpel and steady hand, or a vinyl cutter), but also remains highly scalable with roll-to-roll processing. Second, it features intrinsic bonding capabilities to a wide range of materials that may be required – glass, metals, and plastics. Tape microfluidics have previously been demonstrated in a lab-on-chip setting.^[6,7]^ The requirements for organs-on-chips, however, are higher due to live biological elements as mentioned earlier. Kratz *et al*. only recently presented an excellent characterization of tapes for organ-on-chip use, and demonstrated HUVEC culture in a single-compartment chip.^[8]^ Yet no study to date – including theirs – has implemented a tape-based multi-compartment system, or barrier-on-chip specifically, and demonstrated its biological functionality.

Here, we present the first such study by investigating two candidate tapes and two fabrication methods for their compatibility with epithelial barrier cells. Using commercially available parts and low-cost patterning methods, we design an 8-fold multiplexed barrier-on-chip system with a per-“organ” cost of €0.5 and necessary start-up equipment costs of as low as €100 in an academic setting. We validate the system by demonstrating formation of a tight barrier over 8 days of culture in all “organs”, with good agreement compared to theory as well as barriers grown on gold-standard Transwell permeable supports. We assess biological function in terms of actin and tight junction protein localization as well as metabolic response as a function of permeable membrane pore size in our barriers-on-chips, as well as compared to Transwells. We further investigate stimulation of these small intestine models with capsaicinoids, the active components in chili peppers.

## Results & Discussion

### Device Construction

Our microphysiological system, displayed in Figure 1, features 8 independent “organs”, each with two channels (1.5×0.2 mm2 cross-section) vertically separated by a track-etched membrane. The permeable area of 19±1 mm2 accounts for a ~80% majority of the total cell growth area, important for permeability assays and potential co-culture applications. We construct it solely from tape and polycarbonate (PC) – a plastic well-established as non-cytotoxic and with good tape-bonding characteristics. While the lateral channel geometry can be freely designed with CAD, the channel height is defined by the thickness of the tape. With a tape lamination approach, we can achieve reasonable flexibility also in this dimension.

**Figure 1:**
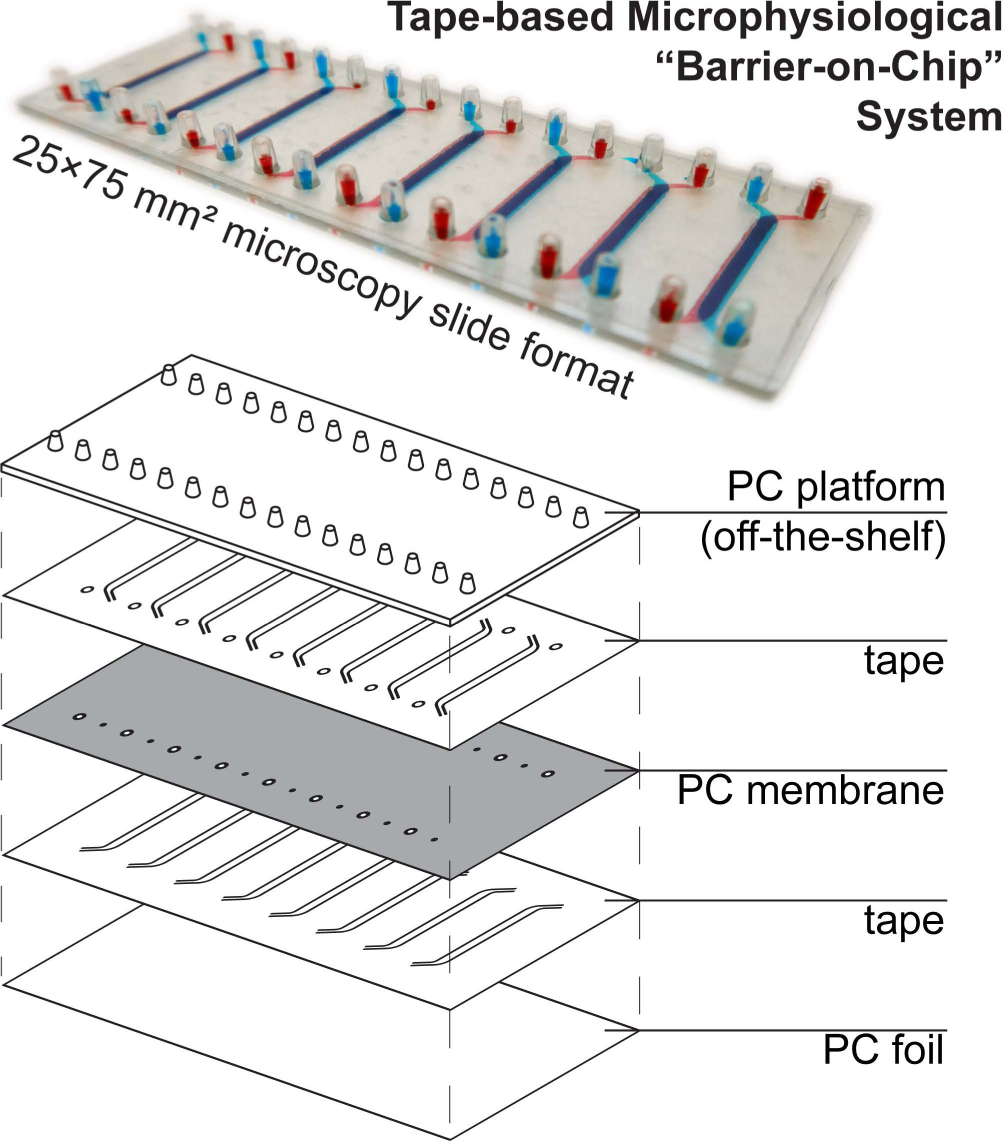
(top) Photograph of assembled device (with intentional top/bottom channel offset for better visualization) perfused with colored solutions. (bottom) Schematic showing device layer structure.

The off-the-shelf PC connector plate accounts for three-quarters of the materials cost budget of our devices. It offers a key advantage over *e.g.* Kratz *et al*.’s use of manually drilled glass slides,^[8]^ however, with a simple chip-to-world interface for elastic tubing (one of the most critical, but also most often overlooked considerations in academic microfluidics). The microscopy slide format also affords simple imaging with existing equipment. For manual assembly, we rely on blunt needle guideposts to provide self-alignment of all layers from the top down, which provides a comparatively rapid and simple process compared to single-“organ” device designs.

### Material & Fabrication Evaluation

Regarding the tapes, we selected two medical-grade candidates for evaluation – the polyester-and-rubber-based 3M 9877, and the polyethylene-and-acrylic-based 3M 9889. They were selected based on high adhesive strength and manufacturer-supplied reports on ISO 10993-compliant cytotoxicity testing. We considered patterning these tapes with either a vinyl cutter (cost from € 100) or a CO_2_ laser cutter (from € 300). The latter has a clear advantage in terms of geometrical freedom over the knife-based approach where sharp corners or small cutouts (< 1 mm) are concerned. The more important consideration, however, is compatibility with the biological elements – in our case, C2BBe1 epithelial cells (a clone of Caco-2 colorectal adenocarcinoma).

In Figure 2, we consider cell coverage after 8 days of culture either directly on top of or adjacent to the two tape candidates, processed with either of the two methods under consideration. We chose to assess coverage rather than viability to eliminate potential interference from loss of dyes for metabolic activity (alamarBlue or similar) to the tape surfaces. We observed the highest coverage (interquartile range (IQR): 98 to 99%) for knife-processed 9877 tape. The acrylic-based 9889 tape shows somewhat more variable growth and/or survival (IQR: 71 to 93%). Laser processing appears to induce generation of cytotoxic compounds especially in the rubber-based 9877 tape, reducing cell coverage by half (95% confidence interval (CI): –20 to –80%) both on and next to the tape.

**Figure 2:**
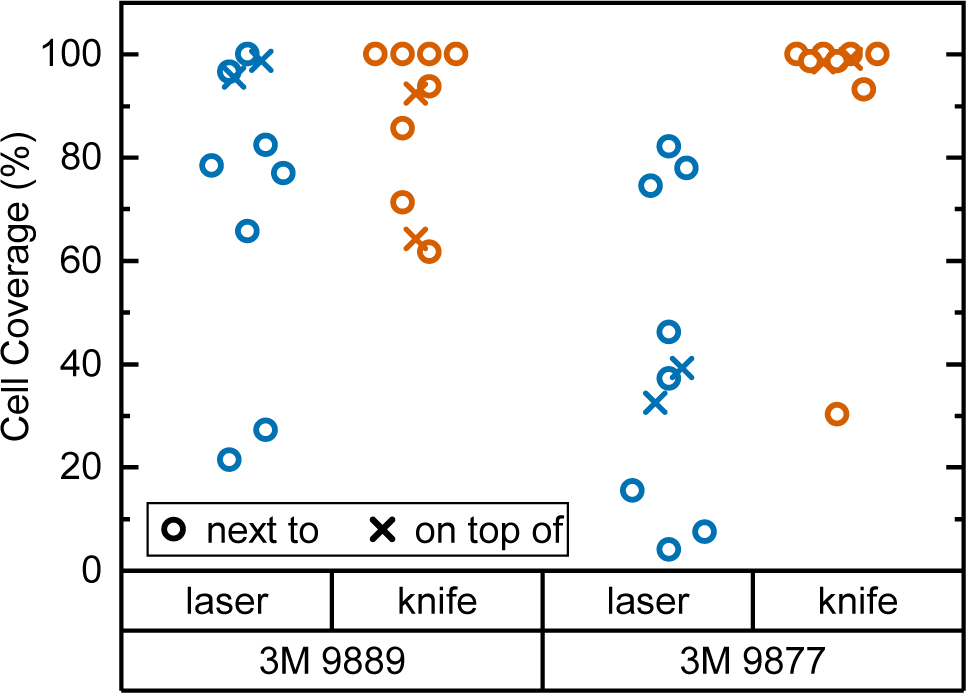
Cell coverage of C2BBe1 cells after 8 days of culture inside a 24-well plate. Well bottoms (N=8) were 50% covered by tape of the type and processing mode given. Cell coverage was assessed both on top of the tape (crosses) and in the non-covered well area next to the plate (circles).

Kratz *et al*., by way of comparison, mainly consider knife-cut ARcare tapes based on acrylic adhesives (akin to 3M 9889). Their cell coverage assessment is qualitatively similar to what we observe with 3M 9877 (rather than the similarly acrylic-based 9889). It may be that ARcare’s acrylics are more suitable for cell contact than the 3M version. However, the different cell type (BeWo b30) and study duration (48h) allows for alternative explanations. Their study did not entail any synthetic rubber-based tapes (akin to 3M 9877).

Another important consideration for microphysiological system construction is the ability to sterilize – or at least disinfect – the materials. The 3M 9877 tape offers an additional advantage in this regard, since it is autoclavable (both 9889 and 9887 can be EtO or gamma-sterilized; we note that ARcare does not provide any relevant information for their tapes). For the assembled systems, however, we relied on disinfection by perfusion with 70% ethanol. The adhesive of both tapes withstands such perfusion for at least 15 minutes, and overall no leakage was observed over 14+ days of perfusion with aqueous solution.

Overall, our initial assessment establishes knife-cut 3M 9877 as a good candidate for organs-on-chips construction based on lack of obvious cytotoxicity as well as its fabrication properties. Our subsequent barrier-on-chip study with C2BBe1 epithelial cells suggests that it is indeed suitable for such applications.

### Biological Validation & Effects of Membrane Pore Size

#### Imaging

We first assess cellular coverage and morphology with fluorescent imaging. This is obtained after 8 days of C2BBe1 cell culture, followed by the 24h capsaicinoid study described in the second part of this paper. In the present section, we will largely limit our analysis to comparisons between the Transwells and tape devices (Figure 3a) that were cultured in parallel. In the widefield images, we observe consistent and complete coverage of all microfluidic channels (Figure 3b) and Transwells (Figure 3c) with epithelial barriers, except for those intentionally disrupted. Some between-channel and within-channel heterogeneity is apparent, but within our expectations for a carcinogenic cell line, as evidenced by the Transwell observations. The intensity variations are moreover partly caused by differing staining efficiency – as with any organs-on-chips, the multiple wash steps in immunocytochemistry present the biggest opportunity for bubble introduction in the overall workflow.

**Figure 3:**
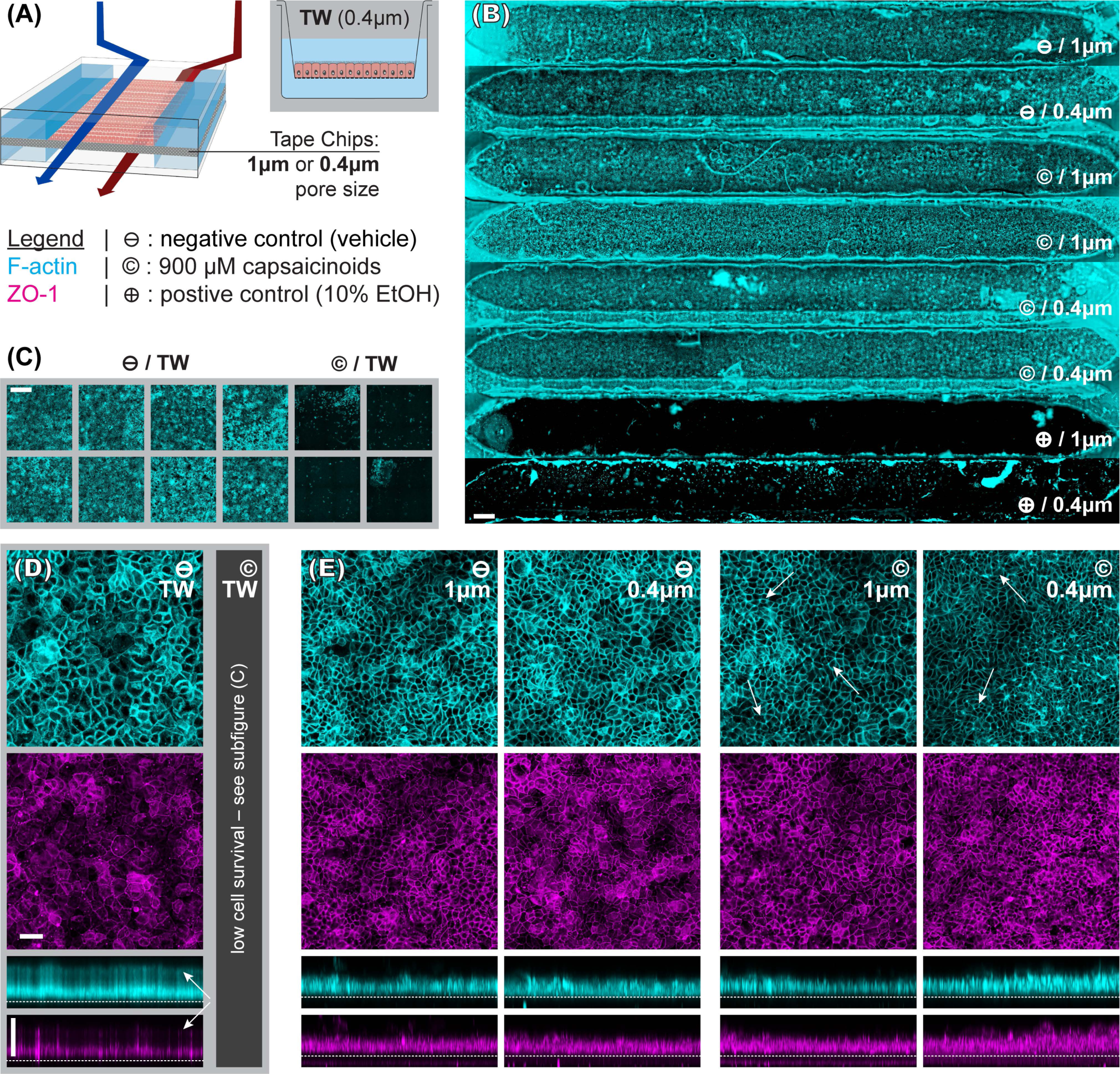
Immunocytochemistry fluorescence micrographs. (A) Overview of the experimental conditions. (B-C) Widefield images (scale bars: 500 μm) respectively illustrating intact epithelial layers across tape chips (⊖ and © conditions, n=6; complete disruption for ⊕, n=2) and Transwells (⊖ condition, n=8; complete disruption for ©, n=4). The fibers are fabrication artifacts. (D-E) Selected maximum intensity confocal projections (scale bars: 50 μm) along z- and y-axes for Transwells (⊖ condition; © was only widefield-imaged due to low survival) and devices (⊖ and © conditions; ⊕ was only widefield-imaged due to low survival). The top surface of the track-etched membrane is indicated with a dotted white line. The white arrows indicate features discussed in the text.

To better compare structural features, the widefield imaging is supplemented by confocal imaging. This reveals significant differences in barrier morphology. Qualitatively, we see more pronounced undulations of the epithelial cell layer in the devices (Figure 3e; y-projections) compared to flatter layers in Transwells (Figure 3d). In the latter, the cell layer is however overall thicker at an average 49 µm (IQR: 47 to 51 µm) compared to 33 µm (IQR: 30 to 36 µm) inside the device channels. Similarly, average cell footprint decreases from 22 µm (95% CI: 21 to 23 µm) diameter to 16 µm (95% CI: 14 to 17 µm) diameter. Thus, cells have similarly columnar shapes in both formats, but are smaller in volume inside our microfluidic channels. It is worth noting here that all the other conditions studied (membrane pore size, capsaicinoids) are associated with well-overlapping morphological IQRs/CIs.

Our observations on cell morphology in devices compared to Transwells are partially in contrast with other works demonstrating thicker epithelial layers in microfluidic culture.^[9,10]^ This may be due to two factors. First, we relied on media recirculation to limit media consumption during maturation (d2-7). However, Shin *et al*. recently found that basally excreted factors in polarized Caco-2 layers can inhibit 3D morphogenesis.^[9]^ In a recirculated system the situation is thus more akin to Transwells, with these factors continuing to reside in the media. Second, our microfluidic channels provide less physical growth height (~200 µm) than some other systems, which has been shown to limit epithelial barrier thickness.^[9]^ The reduced height allows for more even shear distribution (given typical 1-2mm channel width), in our case an average of 0.025 dyne/cm2 (pulsatile; 0.15 dyne/cm2 peak). We note that our epithelial thickness is comparable to other works with similar culture duration, channel height, and shear.^[11]^

Immunocytochemistry reveals one critical difference of biological relevance between static and dynamic culture. In Transwells (Figure 3d, white arrows in y-axis projection), both actin and even more so tight junction protein 1 (ZO1) are almost exclusively confined to the membrane-side. This indicates a lack of (or even inverse) polarization (we will continue to refer to the above-membrane compartment as “apical” in both Transwells and devices for consistency). Microfluidic culture (Figure 3e), on the other hand, shows clear actin expression both basally (membrane adhesion) and apically, where we also observe ZO1 expression. Besides this much-improved polarization in microfluidic culture, tight junction expression is moreover qualitatively more consistent and defined compared to Transwells (z-projections; independent of subsequent capsaicinoid treatment). This is indicative of shear stress in the optimal regime for Caco-2 barrier formation.^[11]^

#### Tracer Permeability

We assess epithelial barrier integrity using Lucifer Yellow (LY) fluorescent dye diffusion. Figure 4 shows that strong barriers have formed by this point in all channels, with apparent permeability coefficients *P*_app_ decreasing by over two orders of magnitude compared to membrane-only controls. After accounting for outliers due to bubbles (asterisk) and those below the assay noise limit, we measure a *P*_app_ of 0.95 nm/s (95% CI: 0.67 to 1.33; averaged over both 1 µm and 0.4 µm membrane pore size) in our tape devices. This is well within the confidence interval of that from Transwells at 0.65 nm/s (95% CI: 0.23 to 1.86; 0.4 µm pore size, but at ~2-fold higher density compared to device membranes). The lower variation within devices compared to Transwells is worth emphasizing, indicating good reproducibility both in device fabrication as well as in barrier formation. Our measured *P*_app_ also agree with Lucifer Yellow literature values for Transwells (Caco-2; 1-7 nm/s)^[12,13]^ or an on-chip model (C2BBe1/HT29-MTX 9:1 co-culture; 1.2 nm/s).^[14]^ While not employing LY, a number of studies have shown similar or even increased *P*_app_ in microfluidic culture compared to Transwells, matching the trends we observe.^[10,15]^

**Figure 4:**
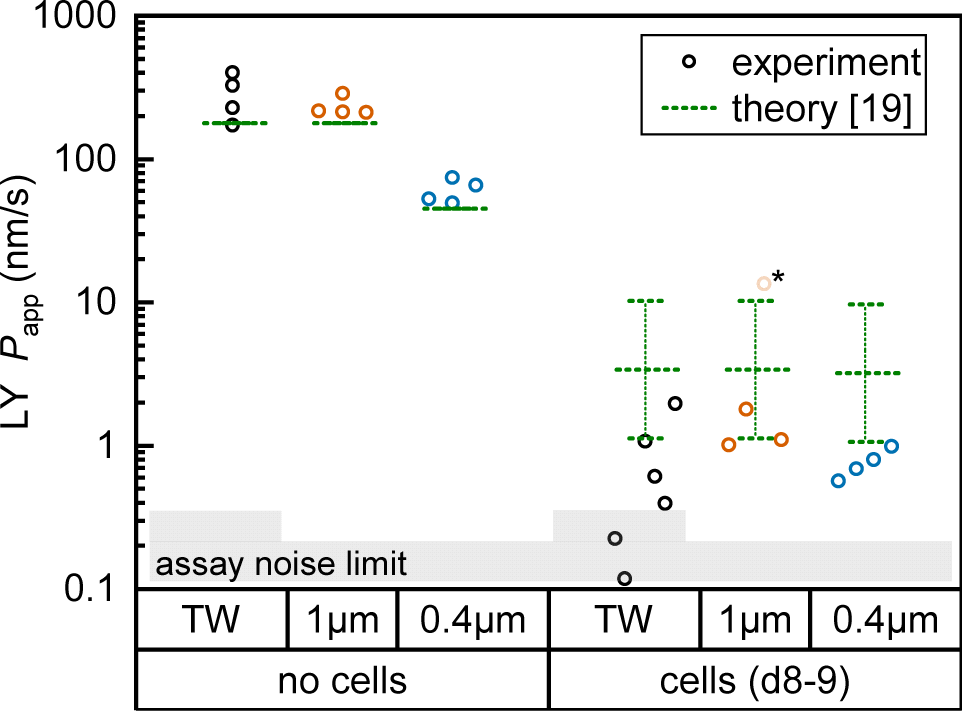
Fluorescent tracer LY permeability compared between Transwells (TW, black) and tape devices featuring either 1 μm (orange; n=4) or 0.4 μm (blue; n=4) pore size membranes. We plot data both without cells (left) and after 8 (TW; n=6) to 9 (on-chip; n=3-4) days of epithelial cell culture. The noise limit for the assay is indicated in gray, higher for TWs due to the less favorable ratio of permeable surface area to basal volume. The starred device datum represents an outlier likely due to a bubble in the channel (*P*_app_ for this device at subsequent timepoints is in line with others). Theoretical predictions for permeability are shown in green.[19] The reference derives an error range by comparing their cell layer model to experimental data for ~150 compounds (the “no cell” condition does not receive similar attention and is not calculated with an error range).

For on-chip culture, we consider whether barrier integrity is dependent on the pore size (0.4 µm or 1 µm diameter) of the track-etched membranes used. We measure an increase in barrier function with decreasing pore size, as *P*_app_ decreases by −0.23 orders of magnitude (95% CI: −0.46 to +0.01 Log_10_) going from 1 µm to 0.4 µm pores. While not previously studied with Caco-2, this aligns favorably with a −0.35 Log_10_ decrease reported for mouse brain endothelial cells in Transwells.^[16]^ Additional Transwell data have been published for human brain endothelium, with mixed trends.^[17,18]^ The study showing higher *P*_app_ with 0.4 µm over 1 µm pores is however confounded by a 2-fold difference in membrane pore densities.

Lastly, we compare measured *P*_app_ values with theoretical ones calculated following the model by Bittermann and Goss.^[19]^ For membrane-only controls, the model consistently underestimates *P*_app_ by ~40%, likely due to errant assumptions about Lucifer Yellow’s diffusion through pores. In the presence of the epithelial barrier (the focus of their work), the model prediction is conversely 3-5× higher than observed values. This is still within the bounds of the predictive power of the model (±0.48 Log_10_), and in line with most of the – like LY^[20]^ – fully ionized species considered in their paper.

#### Metabolomics

Untargeted -omics approaches provide a wealth of biological information. We apply a metabolomics approach to our on-chip and Transwell small-intestine models to gain insight into biological function across conditions. The single-order LC/MS on sampled (for devices, effluent) media captures changes in small (< 1kDa) molecules secreted (and consumed) by the cells. Our study design focuses on global rather than specific changes, *i.e.* considering multivariate and network analysis rather than annotation and validation of a small subset of compounds.

Here, we first take a look at the overall principal component analysis (PCA) across all collected data (Figure 5a). Within two principal components accounting for almost 25% of the overall variation, the unguided clustering proves insightful. PC1 distinguishes well between apical (closed symbols; more positive) and basal (open symbols; more negative) compartments in either culture system. This indicates a sizable conservation in biological function between Transwells and microfluidics. PC2, on the other hand, captures the type of experiment as well as the later-described capsaicinoid condition. Devices (circles) generally score negative, and Transwells (triangles) positive, while capsaicinoid application (red) in either system shows a trend of PC2 towards zero compared to the respective controls (blue). The described patterns are clearest for apical compartments, whereas basal compartments show more overlap between conditions and also with the no-cell media controls. Since basal compartments require permeation of compounds across the membrane, the lower response and closer resemblance to media is expected. Higher-order PCs (not shown) show additional differentiation between capsaicinoid and culture type conditions. The remaining variation among samples is largely due to culture time (causing degradation of compounds in inlet and outlet reservoirs, with additional accumulation effects in Transwells), which is partially reflected in the spread of the media control PCA (green crosses).

**Figure 5:**
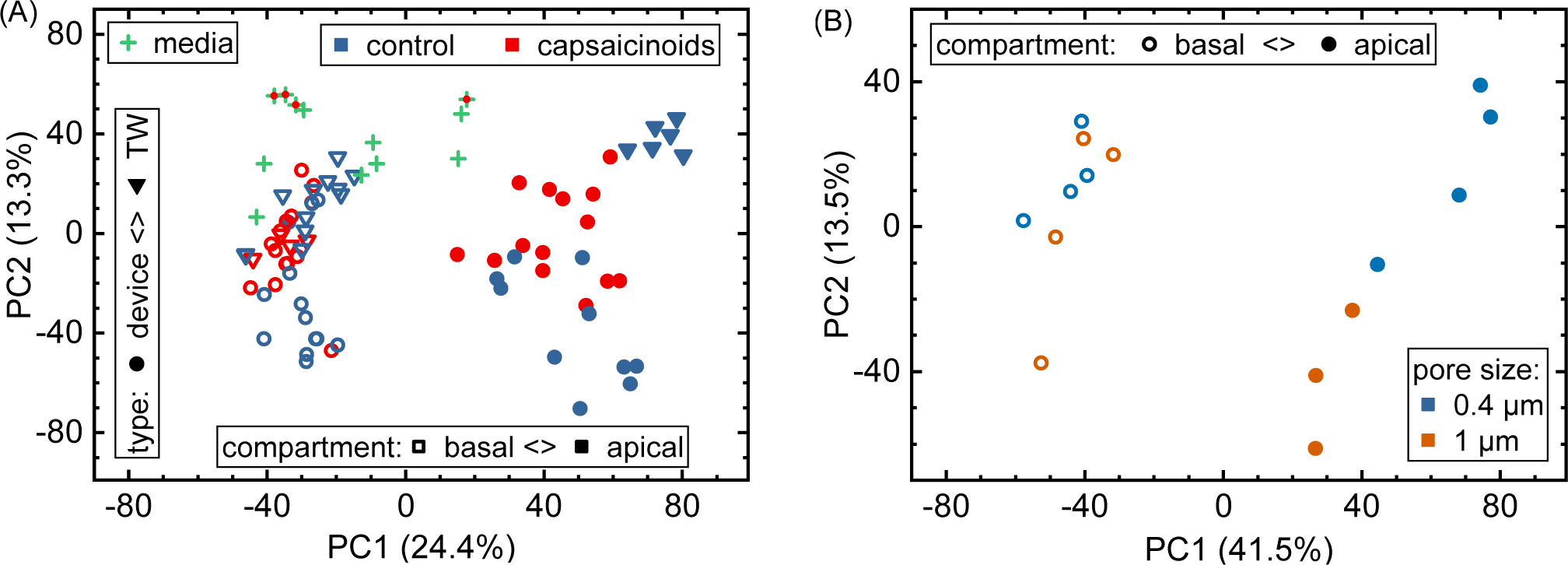
(A) Overall analysis of metabolomic data with PCA. Samples are grouped by culture type (circles *vs.* triangles), apical/basal sampling compartment (closed *vs.* open), and capsaicinoid application (red vs. blue). Media sampled throughout the experiment is also included (green crosses; capsaicinoid media additionally features a red dot). (B) Sub-group PCA showing the effect of permeable support pore size (blue *vs.* orange; N=4 devices each) in our devices. One apical (1 μm) datum is missing due to a sampling issue.

Metabolomic analysis also reveals the clearest differences based on membrane pore size amongst our assays. While not apparent in the full data (Figure 5a), PCA on the relevant sub-set (Figure 5b) shows distinct clusters. Apical *versus* basal sampling again accounts for the bulk of sample variation (PC1). PC2 however differentiates between 0.4 µm diameter pores (blue; more positive) and 1 µm diameter pores (orange; more negative).

Network analysis is more suited for comparisons within the same “organism”, or in our case, culture system. The differing assay kinetics between Transwells (accumulating, and significantly diluting in the basal compartment) compared to devices (continuous perfusion with matched apical & basal volumes) caution against direct comparisons in network activity. One observation that we would like to mention regardless is that Cytochrome P450 activity is among the higher-scoring pathways (when comparing sampled media to blanks) in both Transwells (38 significant metabolites, out of 52 possible within the pathway) and devices (50 significant hits). The extent of this particular pathway in Caco-2 cells is an occasional matter of debate.^[21]^ Our study supports a relatively robust presence in both Transwell and device culture after 8 days, though the nature of our study does not allow for distinguishing enzyme isoforms.

The different membrane pore sizes present a better application for network analysis. While ensemble differences appear in the PCA (Figure 5b), the network analysis does not show any significantly affected pathways (*p*<0.1 cutoff). While differences between pore size are thus present, they appear to be quite subtle, in line also with our other assay results.

### The Effects of Chili Pepper (capsaicinoids)

The above section demonstrates epithelial barrier formation in our tape-based microphysiological systems. We studied the impact of membrane pore size on the cells, and compared our devices to traditional Transwell culture on a number of metrics. To additionally evaluate stimulus response, we select a common food compound: capsaicinoids, the “active ingredient” in chili peppers. They are of potential pharmaceutical interest for pain and body weight management, or cancer treatment.^[22]^ Previous *in-vitro* studies on epithelial barriers have focused mainly on short-term (1-2 hours) cellular response, for a dose range of 100 µM to 500 µM capsaicin applied to Caco-2 or MDCK (a canine kidney epithelial line) barriers.^[23–28]^ These studies have established actin reorganization linked to tight junction opening, and explored the cellular pathways behind it.

#### Capsaicinoid concentration

For our study, we select two nominal apical capsaicinoid levels: 600 µM (the focus for Transwells) and 900 µM (the focus for devices) – the rough equivalent of eating (and efficiently digesting) a very hot habanero chili pepper on an empty stomach.^[29,30]^ While the untargeted single-level LC/MS does not allow for quantification, it does allow us to infer approximate (order-of-magnitude) relations between nominal and effective concentrations. Due to the qualitative nature of this analysis, we will however continue to refer to nominal doses throughout the subsequent sections.

From both Transwell and device data, we can infer an apparent permeability on the order of 100 nm/s for capsaicinoids. In media sampled after 20h of incubation, concentrations are reduced by approximately −0.5 Log_10_ (capsaicin), −1 Log_10_ (dihydrocapsaicin), and −2 Log_10_ (nordihydrocapsaicin). This suggests loss from precipitation (capsaicinoids are poorly soluble in water) and/or slow thermal breakdown.^[31]^ We find that concentrations in devices are an additional −0.5 Log_10_ decreased compared to the media control. We attribute this loss to the microfluidic tubing, which makes up for ~93% of the fluidic surface area. While PDMS can soak up significant quantities of hydrophobic compounds, this is a bulk phenomenon not applicable to our devices, making the tape microfluidics themselves (~6% surface area) the more unlikely candidate.^[32]^ We note that none of the referenced studies on capsaicinoid effects have investigated whether effective concentrations match nominal ones (and in some studies, it remains unclear whether dosing was apical, basal, or global).

Overall, the implications are (1) a rapid apical/basal equilibration in Transwells over short (<4 h) timescales, which we recapitulate in two of our microfluidic channels over the final experimental hours by combined apical and basal dosing; and (2) a lower effective dose in devices, which we seek to compensate for by increasing nominal dose compared to Transwells to the maximum indicated by capsaicinoid solubility and by keeping ethanol vehicle concentration <1%. Our lowered effective doses in devices roughly correspond to eating a habanero chili pepper on a full stomach.^[29,30]^

#### Ethanol vehicle

To rule out effects from ethanol as the vehicle for capsaicinoids, we evaluate Transwells with regular media compared to those spiked with equivalent amounts (0.6% to 0.9%) of ethanol as in the parallel capsaicinoid treatment. These effects prove to be negligible in all analyses. No qualitative morphological differences are observed in imaging. In quantitative TEER and permeability, the group differences score *p*>0.7 and *p*>0.5, respectively. In the metabolomic analysis, at most 1 compound scored *p*<0.05, compared to ≳50 compounds for all other comparisons. We thus treat ethanol vehicle as equivalent to controls in subsequent analysis.

#### Permeability

Over the initial four hours of the experiment, we apply a nominal dose of 600 µM capsaicinoids in the apical compartments. Considering first Transwell TEER (Figure 6a), we observe a slight drop after 30 minutes independent of treatment, indicating simply disturbance of the barrier form the media change. In the subsequent hour, the barrier recovers, with capsaicinoid-treated epithelial layers interestingly showing 30% higher barrier function (95% CI: +5 to +56%) before reverting closer to the controls again by 4 hours. This sinusoidal full-period timeline differs from some reports of a “half-period” reversible drop in TEER.^[27,33]^ A post-exposure TEER increase has been observed with MDCKs – albeit at different timescales – as well as at least once in Caco2 barriers.^[25,27,28]^ Compared to these studies, our sampling intervals may miss out on a larger initial drop due to an overall compressed or expanded time course. More relevantly, the type of “full-period” TEER response we observe has also been reported for natural chili pepper extract on HCT-8 cells (a gut epithelial line like Caco-2).^[34]^ The capsaicinoid mixture in our study is likely more reflective of chili pepper extract than pure capsaicin, though in previous publications the purity of the capsaicin is not always denoted.

**Figure 6:**
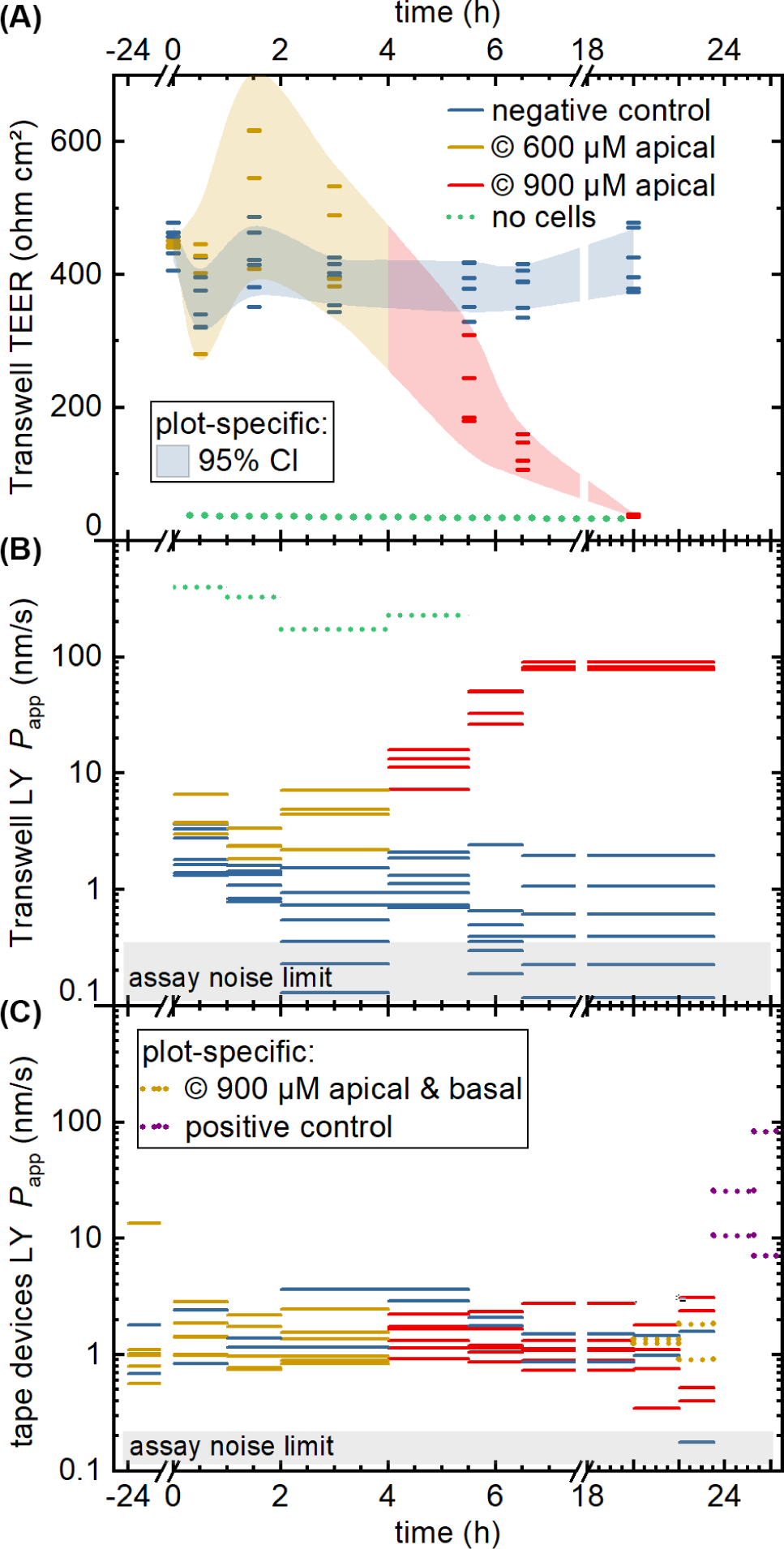
Permeability assays with capsaicinoid application. We compare negative controls (blue) to capsaicinoids at nominal apical concentrations of 600 μm (yellow) and 900 μm (red), as well as a no-cell control (dotted green). (A) Transwell TEER (N=12), with shaded areas tracing the 95% confidence interval. (B) Apparent LY tracer permeability in Transwells, with the assay noise limit indicated by the shaded gray area. Unlike TEER, the permeability assay collects an average over the sampling period, indicated by the length of the lines. (C) Analogous LY tracer assay in microfluidic devices (N=8), with added conditions of global nominal 900 μm capsaicinoids (dotted yellow; n=2) and positive control (dotted purple; 10% ethanol; n=2).

Comparing the TEER now to LY permeability in the Transwells Figure 6b), *P*_app_ similarly shows the negative controls recovering from initial higher values due to the media change disturbance by 2-4 hours. Unlike TEER, capsaicinoid-treated epithelial barriers exhibit consistently worse barrier function in this assay by one order of magnitude (95% CI: +0.5 to +1.6 Log_10_). These opposing trends between assays are intriguing (and were similarly observed in an independent Transwell experiment; data not shown). None of the previously published studies on capsaicin have considered small-molecule tracers alongside TEER. Although LY is generally accepted as a good marker of paracellular permeability, its correlation with gold-standard [^3^H]mannitol is not perfect.[35] Neither is the correlation of TEER with either of these tracer’s *P*_app_. Most intuitive explanations for mismatch apply to a weaker barrier indicated by TEER compared to tracers, due to TEER’s reliance on smaller ions and its relative sensitivity to localized disruption.^[36]^ The case we observe here is more likely to involve transcellular transport. This mode of transport can play a role at TEER values ≳250 Ω cm^2^ if the relevant pores are open at the same time, and can affect *P*_app_ overall.[36] This reasoning would suggest that capsaicinoids can decrease synchronized opening of transcellular ion pores, and/or relatively increase transcellular transport of LY.

By contrast, a subsequent dose increase to 900 µM (4h until endpoint) causes Transwell permeability to increase by both measures (Figure 6a-b). The first 2-3 hours after application already see marked changes, with TEER decreasing by 65% (95% CI: 53 to 76%) and *P*_app_ increasing by 1.8 orders of magnitude (95% CI: +1.3 to +2.3 Log_10_). Unlike the lower-dose effect, this proves irreversible, and both permeability markers approach no-cell values by the next-day experimental endpoint. The initial rapid decrease, and the loss of viability, are consistent with observations of >750 µM capsaicin (apical) or >300 µM (global) in MDCK.^[27,28]^

While the Transwell behavior – in particular the differences observed between TEER and permeability – is intriguing, the comparison to of largest interest to our current study is with tape microfluidic culture (Figure 6c). Here we can only rely on tracer permeability, since a low-cost platform and setup precludes addition of TEER electrodes. As with Transwells, we observe fluctuations even with the negative controls. While the disturbance of cells is less direct than in Transwells, movement of the barriers-on-chips setup in and out of the incubator tends to have larger effects due to lower media volumes (and thus heat retention).

At first glance, the device response in *P*_app_ over time is less pronounced than from the Transwells (Figure 6a *vs.* c). This is in line with expectations due to the different nominal *versus* effective concentrations in the respective systems, as discussed at the start of this section. Nominally, however, the time course mirrors that of the Transwells, at 600 µM apically for the first four hours, and 900 µM thereafter (two on-chip barriers additionally receive 900 µM capsaicinoids basally for the last four hours prior to endpoint). One qualitatively obvious change is when – to assess the validity of the device *P*_app_ assay – we disrupt two “organs” with a cytotoxic dose of ethanol (positive control; 10% apical & basal). This yields an order of magnitude increase in *P*_app_ within the first hour (subsequent readings vary significantly due to dead cells clogging the tubing as verified by visual inspection).

For capsaicinoid application overall, a two-way repeated-measures ANOVA supports a small shift in *P*_app_ either way (95% CI: −0.21 to +0.10 Log_10_) compared to controls. The nominal decrease by −0.06 orders of magnitude appears largely independent of dose, with no sizeable effect even by combined apical and basal capsaicinoid application (900 µM). This stands in contrast to the clear +1.0 Log_10_ increase in *P*_app_ over controls for Transwells at comparable effective dosage and application time (the very initial four hours of the experiment). We suggest two hypotheses. First, the lack of response in microfluidic culture could be due to desensitization from the prior lower dosage. Alternatively, the Transwell culture may overall be more sensitive to capsaicinoid-induced barrier alterations. This type of reduced epithelial toxicity in microfluidic culture has been observed *e.g.* with gentamicin action on MDCK.^[37]^ It is worth noting that *in-vivo* experiments report nominal capsaicinoid concentrations 5-10 times higher than what we observed as cytotoxic in Transwell culture, with minimal adverse effects.^[38,39]^

#### Imaging

Widefield imaging (Figure 3c) reveals nearly complete cell death after the full capsaicinoid time course in the Transwells, in line with the observed TEER and *P*_app_ values. For the devices (Figure 3b), however, the only qualitatively obvious condition is again the cytotoxic dose of ethanol resulting in complete cell death. To discern differences between capsaicinoid and control conditions in the tape microfluidics, we need to consider the confocal images (Figure 3d). Qualitative inspection reveals somewhat less well-defined tight junction expression after capsaicinoid application. We further observe increased actin localization at multi-cell junction points (white arrows). This has been reported for both Caco-2 and MDCK upon capsaicin application.^[26,27]^ The localization is not as common here as in those studies, but ours consider a much longer time course (24h *vs.* 6h). On such timescales, other studies have shown an actin expression profile that mirrors TEER, decreasing initially before increasing significantly over controls.^[24,25]^ This likely results in less organized F-actin repolymerization masking most of the triple-junction localization by 24h.

Quantitative analysis further substantiates alterations in cellular junctions, with actin/ZO-1 colocalization decreasing in terms of Pearson’s *R* from 0.53 (controls) by −0.13 (95% CI: −0.27 to +0.01). This trend is conserved for other correlation measures. A similar analysis grouped by membrane pore size, conversely, shows a negligible change in correlation of 0.02 (95% CI: −0.18 to 0.14). Such colocalization analysis is more robust than direct comparisons of fluorescence intensity. The change in colocalization implies a reorganization of the actin network and/or the cellular tight junctions, in line with our qualitative analysis conclusions.

#### Metabolomics

Our focus for metabolomic analysis is the 600 µM nominal dose for Transwells, and a 900 µM nominal dose for devices. As discussed earlier, this provides us with a comparison of more closely similar effective dosage between culture systems. As also briefly mentioned earlier, PCA of the overall dataset (Figure 5a) shows grouping according to capsaicinoid application compared to controls. The trends (PC2 toward zero) are conserved between Transwells and our tape-based devices, and indeed differences between culture types disappear in the first two PCA components. This suggests capsaicinoid effects dominate over culture type differences, and that biological response is overall quite similar between Transwells and our tape-based barriers-on-chips.

For further insight, we turn to metabolic network analysis (Figure 7). This reveals significant impact of capsaicinoids on the metabolic pathways of our epithelial barriers. Considering first overall changes (Figure 7a), the devices show much more extensive metabolic changes compared to Transwells. We believe this is in part due to dilution effects making detection in the basal Transwell compartment more challenging. However, the two pathways with *p*<0.05 in Transwells are mirrored in devices (*p*<0.1). Two additional high-scoring pathways (*p*<0.05) in devices are further mirrored by the Transwells (*p*<0.1). Biological responses are thus correlated between culture systems not only in PCA, but also on the network level.

**Figure 7:**
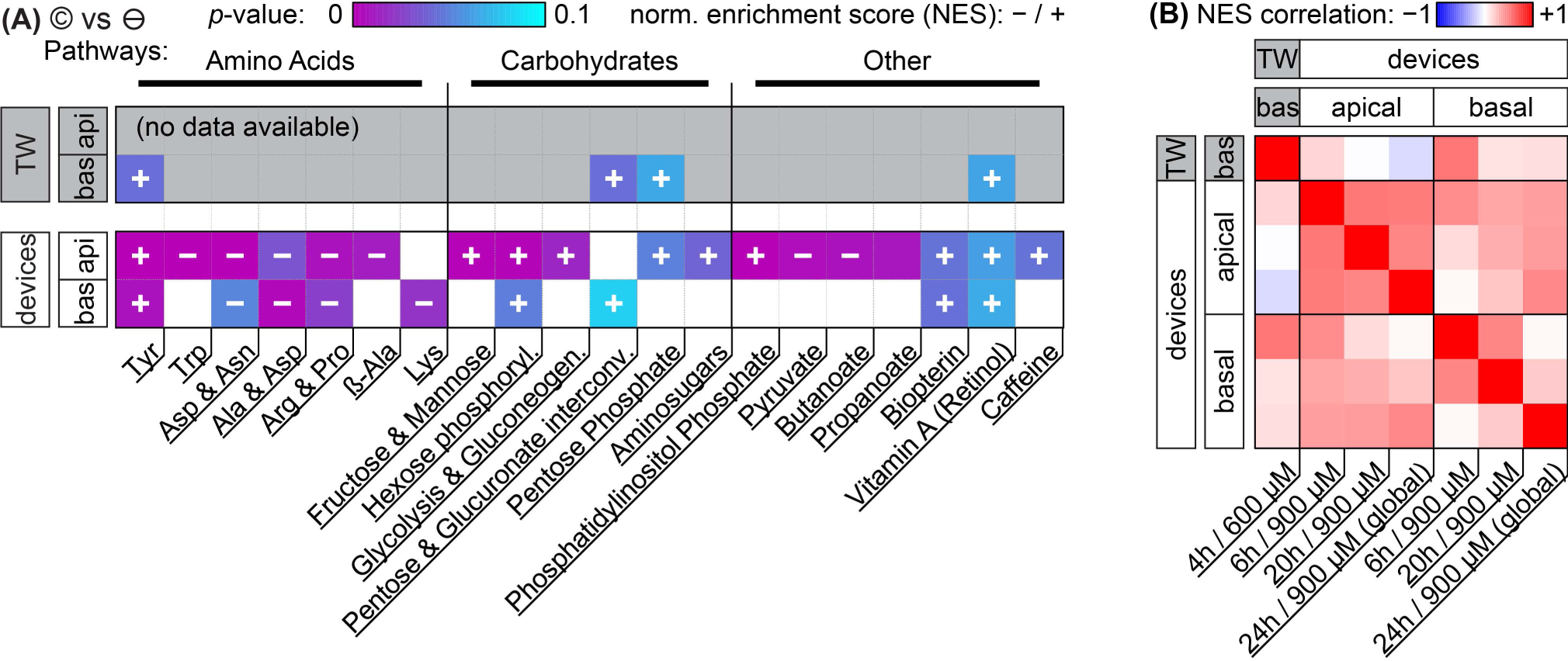
Metabolomic network analysis of capsaicinoid effects (©) on Transwells (TW) and microfluidic devices, separated by basal and apical sampling compartments. (A) The data illustrate significant changes in metabolic networks compared to negative controls (⊖) in terms of *p*-values (color scale from purple to cyan; *p*>0.1 is colored white). Abbreviated network annotations from the MTF metabolic model are listed at the bottom; we have loosely added grouping at the top. The direction of the activity change (in terms of normalized enrichment score NES) is indicated by the overlaid plus/minus signs (missing overlays signify low NES). We include the top 20 pathways (as determined by *p*-values across conditions; smallest pathways < 10 hits excluded). For more closely matched effective dosage, Transwell data (n=4) is for a nominal dose of 600 μm (4h), device data (n=6) for a nominal dose of 900 μm (pooled timepoints). (B) Pearson’s *R* correlation matrix (color scale from anticorrelation in blue to co-correlation in red, with white signifying low correlation) for NES for the different timepoints (Transwells: n=4; devices: n=6, except n=2 for 24h / global dosing) and compartments as labeled on the sides and bottom.

Before considering the specific networks affected, we quantify the correlation in terms of the normalized enrichment (NES) of pathways between Transwells and devices at the various timepoints (Figure 7b). This reveals Transwell (basal) response as “most similar” to devices’ basal compartment at the 6h timepoint (Pearson’s *R* = 0.53; topmost row/leftmost column). Based on effective dosages, we would have expected higher similarity at the 24h timepoint. However, it appears that – in spite of rapid capsaicinoid equilibration in Transwells, and the cells’ weak polarization compared to devices – the Transwell metabolic response remains “polarized”. This matches the conserved grouping along PC1 in PCA with capsaicinoid treatment (Figure 5a). The two globally-dosed devices at 24h, in contrast, show rather apical-like response also in the basal compartment (bottom row/rightmost column). Inherent cell layer polarization thus appears to be less critical for capsaicinoid response than its concentration gradient across the barrier.

The richest amount of information from our assay, however, is contained within the metabolic pathways affected in devices (Figure 7a). We broadly group the pathways into “amino acid” metabolism, “carbohydrate” metabolism, and “others”. Tyrosine metabolism is one of the strongest hits overall and upregulated in every compartment and timepoint (including Transwells), while other identified amino acid pathways are downregulated. Most of these pathways are sizable and not straightforward to interpret. The tyrosine metabolism, for instance, involves dopamine and adrenaline. These are consistent with stress modulation as discussed later (for “other” pathways).^[40]^ The literature also provides a more capsaicinoid-specific line of reasoning. Capsaicin happens to be a structural analog for tyrosine, and can compete with the amino acid in its tRNA aminoacylation.^[41]^ This would leave excess tyrosine available for other cellular pathways, consistent with the increased tyrosine metabolism we observe.

Besides amino acid pathways, various networks involved in carbohydrate meta- and catabolism are indicated and generally upregulated. This appears to be in agreement with findings of increased energy metabolism in a range of *in-vivo* studies.^[42]^ For Caco-2 cells specifically, one prior study also describes increased energy metabolism along with overexpression of two particular enzymes involved in glycolysis.^[43]^ Our network-level approach is less sensitive to changes in individual enzymes, but the glycolysis pathway involving these enzymes is present, and the other upregulated “carbohydrate” pathways identified are generally closely adjacent.

The “other” category groups a range of pathways not so easily categorized. It includes small-molecule pathways (pyruvate, biopterin, …) which are closely intertwined with the energetic and stress responses of other networks. In terms of stress and inflammation, the retinol pathway deserves separate attention. Increases in gut retinoid metabolism have been documented for a range of stressors in rats.^[44,45]^

Phosphatidylinositol phosphate is similarly indicated in stress response.^[46]^ Even more relevantly, this pathway is closely linked to cytoskeletal reorganization, aligning well with our (and others’, as discussed earlier) imaging results.

We are aware of one further study considering metabolic effects of capsaicinoids on Caco-2 barriers.^[47]^ Rohm *et al*., using a targeted (and thus more individually sensitive) approach find increased acetyl-coenzyme A synthetase activity, indicating higher fatty acid biosynthesis. For our untargeted assay, however, the overall number of compound hits within this pathway is comparatively low (<30% coverage; compared to ≳50% for the pathways we report on), providing a likely explanation for its absence in our analysis.

Overall, we thus find a complex metabolic response to capsaicinoids in our on-chip barriers that aligns with other *in-vivo* and *in-vitro* findings from literature. This high metabolic response – sustained through all timepoints (Figure 7b) – combined with (compared to Transwells for similar effective dose) much less pronounced changes in barrier integrity presents an argument against the desensitization hypothesis we offered in the earlier analysis (*cf.* permeability discussion). Instead, the response we observe in our devices may indeed reflect a more physiological-like response to chili peppers.

## Conclusions

Our approach to barrier-on-chip system fabrication based on double-sided pressure-sensitive adhesive tape proves to be both affordable and functional. For small intestine modeling, we obtain an epithelial barrier layer with tight cellular junctions across the entirety of the microfluidic channels. Cell polarization, barrier permeability, and junction protein expression are consistent with prior *in-vitro* research both in Transwells and in traditional PDMS-based organs-on-chips. We further demonstrate biological response to chili peppers (capsaicinoids). Though devices and Transwells show different response in terms of permeability, our observations on metabolic impact as well as actin and tight junction protein localization align well with other *in-vivo* and *in-vitro* studies.

Our tape-based approach to barriers-on-chip cannot compete with PDMS devices in terms of design freedom, and it remains to be seen whether the materials would be compatible with more sensitive cellular models (*e.g.* hiPSC-derived). Instead we provide a simple and accessible fabrication approach suitable for work with similarly (compared to *e.g.* hiPSC culture) much more affordable cell culture. Tape-based barriers-on-chips can be assembled in academic labs with minimal equipment cost. The well-developed film- and tape-converting processes available, combined with pick-and-place technology, should further make industrial manufacture feasible at much lower per-unit cost than existing systems. We therefore hope that tape-based barriers-on-chips will enable more labs in lower-resource environments to enter a field currently dominated by relatively few well-funded facilities.

## Acknowledgements

We appreciate the support of Dr. Charles Vidoudez and the Harvard Center for Mass Spectrometry in the metabolomic analysis. We further appreciate the assistance of the staff at the Karolinska Institute Biomedicum Imaging Core. T.E.W. is grateful for funding from the European Union’s Horizon 2020 Research and Innovation Program under the Marie Sklodowska-Curie grant agreement “NeuroVU” (No. 797777). M.F. and E.F.G.J.S. acknowledge support from the Erasmus+ program of the European Union. A.H. acknowledges funding from the Knut and Alice Wallenberg Foundation, Göran Gustafsson Stiftelse, and Forska utan Djurförsök.

## Materials & Methods

### Organ-on-a-Tape-Chip Fabrication

Double-sided medical-grade adhesive tapes types 9889 and 9877 for material evaluation was kindly provided by 3M (Maplewood, Minnesota). Additional 9877 tape for device manufacture was obtained from Beneli AB (Helsingborg, Sweden). 9877 tape has a total thickness of 110 µm, with a synthetic rubber-based adhesive and a 23 µm polyester film carrier. 9889 tape has a total thickness of 120 µm, with a tackified acrylic adhesive and a 80 µm polyethylene carrier. Polycarbonate (PC) membranes (25 µm thick) with track-etched pores (1.6×10^6^ cm^−2^, 0.4 µm or 1 µm diameter) were purchased from ip4it (Louvain-la-Neuve, Belgium). 125 µm thick PC foil (Makrofol DE 1-1) was kindly provided by Covestro AG (Leverkusen, Germany).

We employed a cutting plotter (CE 5000; Graphtec, Tokyo, Japan) and CO2 laser (VLS 2.3; Universal Laser Systems, Scottsdale) to pattern the tape and membrane based on CAD drawings. Bonding between layers was facilitated using a hydraulic press (Rosin Tech Products, Los Angeles, CA).

### Cell Culture

Human enterocytes (C2BBe1; a clone of Caco-2) were obtained from ATCC at passage 47. Frozen stocks were expanded according to supplier protocols and maintained at 37 °C / 5% CO_2_. Cells were used at passages 50-55. Media was prepared from DMEM (high glucose; Gibco 10569010) and 100 U/ml penicillin-streptomycin (Gibco 15140122). For the cytotoxicity assay, we supplemented the media with 20% heat-inactivated fetal bovine serum (FBS; Gibco A3840002). In the remainder of our study, we employed 10% FBS combined with 1× Insulin-Transferrin-Selenium (ITS; Gibco 41400045). Media was prepared the day prior to use and placed in the incubator overnight to equilibrate (critical to avoid bubble formation in the tape microfluidics).

### Cytotoxicity Testing

The cell coverage experiments were conducted in 24-well plates (flat bottom, TC-treated). Semicircular pieces of tapes were cut and inserted into the wells (N=8). In this experiment only, no attachment-supporting coatings were applied, but we relied on proteins adsorbing from the high 20% serum content in the media. Cells were seeded out over the entire well area and cultured over eight days. We manually segmented phase contrast images to estimate cell coverage next to the tapes (n=4 images per well). The coverage analysis for growth on top of the tapes relied on threshold segmentation of the Hoechst fluorescence signal (n=1 image per well). Each image/datum corresponds to one 9.5 mm2 field of view (10% of the total available cell growth area).

### Experimental Procedure

For the biological functionality study, we conducted parallel experiments with tape microfluidics (N=8; n=4 per membrane type) and Transwell Permeable Supports (N=13; n=1 as no-cell control) with a 10 µm, 0.33 cm^2^ polyester membrane (0.4 µm diameter pores; 4×10^6^ cm^−2^; Corning 3470). Transwells were coated with a mixture (prepared on ice) of 300 µg/ml Matrigel Growth Factor Reduced (GFR) Basement Membrane Matrix (Corning 354230) and 50 µg/ml Collagen I Rat Tail High Concentration (Corning 354249) to enhance cell attachment by overnight incubation at 37 °C. After removing excess coating solution, enterocytes were seeded at 1.5×10^5^ cm^−2^ on top of the permeable supports along with 200 µl media, and 800 µl media in the bottom (basal) compartments. Media was exchanged every second day until day 8 (only half the media volume was replaced in the apical compartment).

For tape microfluidic culture, flow was provided by a 16-channel peristaltic pump (ISM 1136; Cole-Parmer, Wertheim, Germany) featuring 0.25 mm inner diameter (ID) PharMed BPT tubing. An additional 5 cm of the same tubing provided chip access on either side, and longer sections were used when recirculating media. Media reservoirs consisted of sterile-packaged 6 ml syringe bodies with blunt needles and 12 cm of 1.6 mm ID Tygon ND-100-65 tubing (Saint-Gobain, La Défense, France). Fluidic interconnects were fashioned from 1.6 mm ID tubing or 0.2 mm ID stainless steel pins (Interalloy, Schinznach-Bad, Switzerland) as needed. Blunt needles, pins, and tubing were autoclaved before use. The pump was controlled via a custom LabView interface to generally provide an average forward flow of Q~90 µl/h. The specific pulsed scheme applied a combination of: paused / 540 µl/h forward / 540 µl/h backward / 2700 µl/h flush at a duty cycle of 50% / 33% / 16% / <1%.

On day −1, devices were disinfected (and cleared of bubbles) by flushing with 70% ethanol for 5 minutes. Subsequently, they were generously rinsed with PBS, again ensuring no bubbles remained in the channels. We then aspirated a collagen/matrigel coating (50 µg/ml & 300 µg/ml) into the channels via pre-chilled tubing. Cell culture media reservoirs were connected, but perfusion was only started after an initial static incubation for 6 h at 37°C. On day 0, enterocytes were seeded in the upper channel at 1.75×10^5^ cm^−2^ via aspiration and left to attach in the incubator under static conditions for 2 hours, before re-starting perfusion. We gradually ramped up the flow to 90 µl/h over 12 hours before switching to the aforementioned pulsed scheme. The cells were allowed to form a barrier over 8 days, with media recirculating from day 2. On day 7, recirculation was stopped and fresh media introduced.

On day 8, media was fully exchanged in all Transwell compartments and microfluidic reservoirs (including inlet connection tubing). Basally, we supplied regular media in all conditions. Apically, we supplied either regular media (n=4 Transwells), media containing 600 µM capsaicinoids (n=4 Transwells, n=6 on-chip barriers), or media containing an equivalent amount of ethanol vehicle (0.6%; n=3 Transwells, n=2 on-chip). Conditions were distributed equally among membrane types for the devices. Capsaicinoid media was prepared by spiking from a 100 mM ethanolic stock solution 30-60 minutes prior to use, followed by gentle agitation and returning it to the incubator until needed. Dosages were increased to 900 µM (or 0.9% ethanol vehicle) after four hours by additional spiking into apical Transwell compartments or device inlet reservoirs, making sure to empty out device tubing of lower-concentration media. Negative control microfluidics received similar treatment to ensure consistent disturbance of the cells from handling. At twenty hours, n=2 on-chip cultures additionally received 900 µM capsaicinoids basally, and were changed again to positive control (10% ethanol) after 210-240 minutes.

### Permeability Assay

Transwell TEER was measured with an EVOM2 instrument and STX2 chopstick electrodes (World Precision Instruments, Sarasota, Florida). Due to inadvertent media cross-contamination between two Transwells (negative control & vehicle), these are excluded from all permeability and metabolomic analysis.

For tracer dye permeability we relied on Lucifer Yellow (Thermo Fisher L453) added to the media at 1 mM in the apical Transwell compartment or apical microfluidic channel inlet. To facilitate concentration equilibration in the basal Transwell compartments, the cultures were placed on a rocking plate inside the incubator.

Basal Transwell compartment or basal microfluidic channel outlet samples of *V*_sample_=50 µl were collected at each timepoint *n*, transferred into a 96-well half-area flat bottom microplate, and measured using an Infinite 200 Pro fluorescence reader (λ_ex/em_=418/475 nm; Tecan, Männedorf, Switzerland). Serial dilution curves of reference (inlet) media were fitted with a Hill function to calculate fractional dye concentration *C* in basal (outlet) samples for each timepoint. Permeability in the devices was calculated as

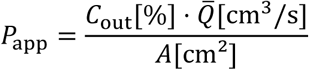

where *Q* is the average volumetric flow rate, and *A* the permeable membrane area. For Transwells, the formula is more complex due to dye accumulation over time *t*. We moreover need to account for successive dilution from fresh media addition after each sampling step to keep the basal volume constant at *V*_basal_=800 µl.

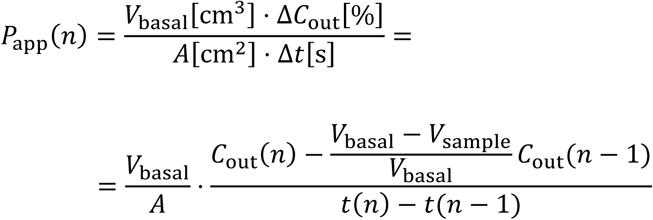

### Immunostaining

For Transwells, media was aspirated and cells washed with PBS. Cells were fixed with 4% paraformaldehyde for 20 minutes at room temperature. *Caution: paraformaldehyde may cause cancer and is suspected of causing genetic defects. Handle with care and proper protective equipment in a properly vented environment.* After three washes with PBS, 50 µl blocking buffer (10% goat serum (Merck G9023) and 0.1% Triton X-100 in PBS) was added to each well and left to incubate for one hour at room temperate on a rocking plate. Subsequently, we washed once with PBS and added 50 µl primary antibody solution in each well. This solution (10% blocking buffer in PBS) contained mouse anti-ZO1 antibody (Invitrogen 33-9100) at 2.5 µg/ml and was left in the wells overnight at 4°C. The next day, after two PBS washes, we added 50 µl secondary antibody solution containing 10 µg/ml goat anti-rabbit CF594 (Sigma SAB4600107) and left this to incubate one hour at room temperature. Finally, after two PBS washes, we added 50 µl of the final counterstaining solution and incubated for 30 minutes at room temperature. This solution (10% blocking buffer in PBS) contained 4 µM Hoechst 33342 (Invitrogen H3570) and 6.6 µM phalloidin AF488 (Invitrogen A12379). Devices were washed three times with PBS and stored in the dark at 4°C until imaging. Prior to imaging, we excised the Transwell membranes using a scalpel and mounted them using Vectashield antifade mounting medium (Vector Labs H-1000) between glass slides.

Device immunostaining proceeded with the same solutions and incubation times. Solutions were aspirated using the pump, and for incubations < 1h agitated by intermittent flow. Washing was replaced with ~10 minutes PBS perfusion for each wash step.

### Imaging

We acquired widefield fluorescent images of all N=8 tape microfluidic channels and N=12 Transwells. Additionally, we selected n=2 spots per device and n=1 spot per Transwell for confocal imaging. Spots were chosen near either end of the devices along the centerline, and near the center of the Transwells. We avoided areas with obvious imaging impediments (*e.g.* the fibers evident in Figure 2), but otherwise made no attempts to optimize frame selection. For Figure 3d-e, we randomly selected one of the two spots per device, and two representative images from the Transwells.

We relied on Fiji for all image analysis.^[48]^ Confocal images were de-noised with the CANDLE algorithm,^[49]^ and corrected for xy-plane tilt using TransformJ. Maximum intensity projections were calculated along x-, y-, and z-axes. In our microfluidic devices, the membrane (and its pores) yielded a signal comparable to that from cellular structures, likely due to inefficient blocking prior to antibody introduction. The z-projection calculations thus exclude the relevant z-slices. All fluorescent images were processed with CLAHE to emphasize local structural differences over longer-distance intensity variations which at least in part arise from staining variations within and across the microfluidic channels.

To estimate layer thickness, we applied auto-threshold segmentation to the x- and y-axis projections. The extracted thickness histogram was fitted with a Log-Normal distribution. For correlation analysis, we relied on the Coloc plugin to extract Pearson’s *R* and related measures.

### Mass Spectrometry Analysis

Samples for metabolomic analysis were collected by freezing down permeability assay samples immediately after fluorescence measurement, and storing them at −80°C. A total of N=73 samples were selected for analysis, encompassing timepoints 0h, 4h, 6h, and endpoint for devices, and timepoints 4h and endpoint for Transwells. The selection favored later timepoints to increase metabolite accumulation, but excluded Transwell timepoints where cell death was inferred for the higher capsaicinoid dosage (see Results section). Additionally, N=11 cell culture samples from device inlets or the no-cell Transwell were included at various timepoints (including fresh media, and a freshly-prepared dilution series of capsaicinoids to estimate dosage). N=11 quality control samples were prepared by pooling aliquots from the 73 assay samples. The dual-polarity LC-MS on an Orbitrap ID-X (Thermo Fisher) coupled to a ZIC-pHILIC column (2.1×150 mm, 5 µm; Millipore, Billerica, MA) was performed as previously described.^[50]^

The raw data were converted into an open-source format using ProteoWizard.^[51]^ We employed XCMS online for feature alignment,^[52]^ with parameters derived from quality control samples using IPO.^[53]^ The data and XCMS analysis are available online.^[54]^ This pipeline yielded ~18,000 features. Drift compensation was performed with statTarget based on the quality control samples (QC-RFSC algorithm), after filling missing features using half-minimum estimation (within-group 70% rule).^[55]^ We further discarded features with remaining quality control RSD > 40%. Lastly, we manually filtered out features associated with capsaicinoids themselves. We relied on a combination of the following criteria: known isotopes/adducts of capsaicinoids; grouping classification by XCMS CAMERA; increasing feature intensity with concentration in the capsaicinoid pseudo-calibration samples; and >5-fold feature intensity in capsaicinoid references over regular media controls. This left us with ~12,500 features for analysis. Univariate and multivariate analysis were performed with statTarget (data were generalized log-transformed and mean-centered). The between-groups FDR-corrected *p*-values and associated fold-changes were used as the inputs for network analysis. Therein we relied on MetaboAnalyst, specifically its combined mummichog (*p*<0.1 cutoff) + GSEA approach using the MTF metabolomic model (5 ppm mass accuracy).^[56]^ We discovered that the GSEA component of this analysis pipeline proved unstable upon repeat analysis. Therefore, we combined GSEA *p*-values and NES scores from 9 analysis runs by geometric^[57]^ and arithmetic averaging, respectively, before integrating this with the mummichog results through Fisher’s method as intended by the authors.^[58]^

